# Bovine brucellosis: seroprevalence, risk factors and assessment of knowledge, attitude, and practice of cattle owners in Lare and Jikawo districts of Gambella region, Ethiopia

**DOI:** 10.1101/2023.01.26.525782

**Authors:** Tamirat Zelalem Kumsa, Bizunesh Mideksa Borena, Biniam Tadesse Derib, Abebe Garoma Gichile, Lencho Megersa Marami

## Abstract

**Background:** Bovine brucellosis is a zoonotic disease that causes substantial economic losses and strongly impacts public health. Though it has been eradicated in many developed countries, it is still endemic in developing countries like Ethiopia.

**Methodology/principal findings:** Lare and Jikawo were the two districts of the Gambella Region selected purposively. Kebeles, study animals and peasant associations were randomly chosen. A total of 384 serum samples from 70 herds were collected and screened using Rose Bengal Plate Test (RBPT) and confirmed using Complement Fixation Test (CFT). A semi-structured questionnaire survey was used. The seroprevalence of brucellosis was summarized using descriptive statistics, and the association between risk factors, and seroprevalence of brucellosis was evaluated using logistic regression. The principal findings of the current study showed that individual and herd level seroprevalence of brucellosis using RBPT was 6.77% (26/384) and 24.29% (17/70), respectively, and the respective confirmation by CFT 3.13% (12/384) and 12.85% (9/70). Among the risk factors, herd size and the presence of other species had statistically significant associations (p<0.05) with Brucella seropositivity. Although the overall respondents’ knowledge, attitude, and practice were 66.4%, most were unaware that the disease was zoonotic, the ability of the disease to cause abortion, and the mode of the disease’s transmission. Most respondents also had a poor attitude toward the mode of disease transmission, and they have been practicing risky practices that predisposed them to brucellosis.

**Conclusion:** The overall seroprevalence of brucellosis and cattle owners’ knowledge, attitude, and practice in the current study were low. However, being a contagious disease, brucellosis can easily spread among cattle herds, and poses a public health risk., Therefore, improvement of cattle owners’ knowledge, attitude, and practice and characterization of circulating Brucella species in the study areas are needed to design evidence-based disease control measures.

**Author Summary:** Bovine brucellosis, is a bacterial disease caused by *Brucella abortus*, which primarily affect cattle. Although bovine brucellosis has been eradicated in many developed countries, it is endemic in Ethiopia. It is one of the most serious zoonotic diseases widely distributed and resulted in serious economic losses and public health impacts. Particularly in pastoral parts of Ethiopia, like the current study areas, factor such as limited veterinary services and education services, frequent movement of farmers from one place to another in search of feed and water for their animals facilitates disease transmission between animals and to human. Thus, it is necessary to assess the prevalence of the disease and awareness of the farmers about the disease in order to reinforce the existing disease control attempts in the country and reduce public risk. Serum from blood samples of cattle were tested using Rose Bengal Plate Test and Complement Fixation Test to detect the presence of *Brucella* antibody, which indicates the presence of bovine brucellosis in the areas. Moreover, farmers’ knowledge and practice regarding the disease information was assessed. The current study showed the presence of brucellosis and farmers had low knowledge, attitude and practice risky activities that predispose them to the disease.

## Introduction

Brucellosis has been eradicated in many developed countries; however it is still endemic in developing countries because of a lack of control programs and/or resources [1]. It is caused by Brucella species, gram-negative, facultative intracellular bacteria that can infect many species of animals. The disease has been reported in many countries around the world, including Ethiopia [2, [3]. In cattle, brucellosis is predominantly caused by *B. abortus*, less frequently by *B. melitensis* and occasionally by *B. suis* [4].Direct contact with infected abortion materials, inhalation, and the consumption of infected milk and milk products are significant means of transmission of the disease to humans [5]. However, infection through injured/intact skin, the mucosa of the respiratory system, and conjunctiva occur frequently [6]. Transmission to animals occurs mainly by ingestion of contaminated feed and wate [7].

Brucellosis is endemic in most African countries [8]. It is considered to be an occupational disease that mainly affects abattoir workers, farm laborers, animal keepers, butchers, veterinarians and laboratory workers from a public health point of view [9]. However, abattoir workers are more prone to acquire brucellosis than other occupations, because they are more exposed to carcasses, viscera, and organs of infected animals [10]. The economic significance of brucellosis results from production losses associated with abortions, retained placenta, metritis, impaired fertility and arthritis. Milk production losses in infected dairy cows can be up to 20% and the inter-calving period can be prolonged by several months [8].

The risk factors that influence the spread and maintenance of brucellosis are related to management systems, the genetic content of susceptible animal populations, the biology of agents causing the disease, and environmental factors. These factors include the size and composition of the herd, age of the animals, frequent contact between infected and susceptible herds, poor farm biosecurity and climate change, abortion, number of parity and age at animal population level [11, [12].

Various serological tests have been developed and are being used to provide rapid results [13]. The standard Rose Bengal and Complement Fixation tests are the main serological tests used to detect antibodies against *B. abortus* and *B. melitensis*. Both tests have been used for several years for eradication of bovine brucellosis in some countries [14]. Different authors have reported evidence of *Brucella* infection in Ethiopian cattle using various serological tests Accordingly, relatively high seroprevalence of brucellosis (above 10%) has been reported from smallholder dairy farms in central Ethiopia [6] while most of the studies suggested a low seroprevalence (below 5%) in cattle under crop-livestock mixed farming [15, [16]. Asmare et al. [17] reported a pooled national estimate of brucellosis of dairy cattle in Ethiopia was 3.3% and Tadesse [18] also reported 2.9%.

Brucellosis is a zoonotic disease that leads to considerable morbidity [19]. The economic and public health impact of brucellosis remains a concern in developing countries [20]. It is among the top five priority zoonotic diseases in Ethiopia [21]. In pastoral societies, brucellosis constitutes significant public health importance where close intimacy with animals, raw milk consumption and low awareness of zoonotic diseases facilitate its transmission between livestock and humans. More importantly, traditional management systems of pastoral communities, such as communal grazing, importing animals from infected herds, intermixing their livestock at water points and using single bulls for breeding purposes without testing, indicate the need for the study of brucellosis in pastoral communities. There is no published literature about the prevalence of cattle brucellosis, level of awareness of cattle owners about brucellosis, and risk factors for the occurrence of brucellosis in Gambella Region, Ethiopia. Therefore, the present study was aimed to estimate the seroprevalence of bovine brucellosis, identify its risk factors and assess the knowledge, attitude, and practice of cattle owners in selected districts of the Gambella region, Ethiopia.

## Material and methods

### Description of the study area

A study was conducted in Gambella National Regional State of Ethiopia, situated in the Southwestern part of Ethiopia, with a total land area of 25,521 square kilometers. It shares a long border with South Sudan and two other Ethiopian regions: Oromia to the North and East and the Southern Nations, Nationalities and Peoples’ Regional State to the South [22].

Jikawo district is one of the four districts in Nuer Zone, located 120 km from Gambella town. It is bordered on the south by Anuak Zone, on the west by the Alwero river which separates it from Wentawo, on North by the Baro river which separates it from South Sudan, and on the East by Lare. The terrain consists of marshy and grasslands with an annual average rainfall of 870 millimeters. It has an altitude of 320-500 meters above sea level and the major crops grown are Maize and Sorghum. The majority of the community are agropastoral and pastoralist and the total cattle population of the districts is 38,834 [23].

Lare district has an altitude of 410-430 meters above sea level and it is located 45 km from the center of Gambella town near the South Sudan border of the Gambella Region of Ethiopia. It is part of the Nuer Zone; Lare is bordered on the South and East by the Anuak Zone, on the west by the Baro River separating it from Jikawo, and on the north by the Jikawo River separating it from South Sudan. The landscape consists of marshes and grass lands with an annual rain fall of 1900-2100 millimeters. The main crops growing in the district are corn, maize, sweet potato, and peanuts. The livelihood activities of the community are pastoral and agropastoralism and the cattle population of the area is estimated to be 45,540. The majority of animals are managed under extensive system by smallholder [24]. The location of study areas is shown on the map (Fig 1).

**Figure 1.**
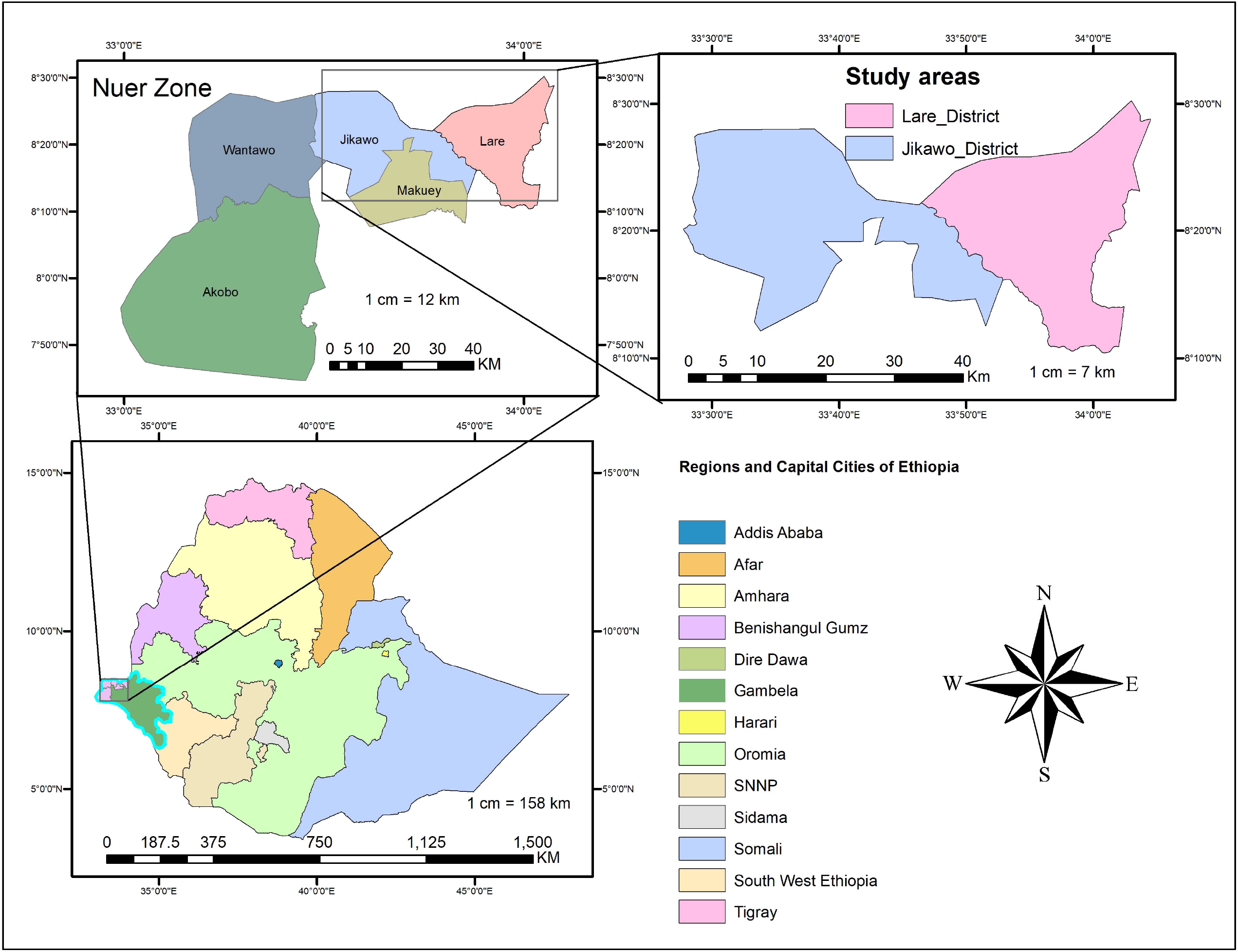
Map of study areas (ArcGIS 10.2.0.3.3348 ESRI).

### Study design and study population

A cross-sectional study, consisting of a questionnaire survey and serology tests, was conducted from October 2019 to April 2020 in the Lare and Jikawo districts of the Nuer Zone of the Gambella region, Southwest Ethiopia. All cattle found in Lare and Jikawo districts were considered as study populations. There are 24 kebeles in the Lare district and 22 kebeles in Jikawo. Kebele is the smallest administrative unit in the district. Ten kebele from each were chosen based on their proximity to transportation. Two kebeles from each of the ten kebeles that are near transportation were chosen at random using a lottery technique. The target populations were cattle (both male and female), over six months of age and reared under an extensive management system in the study areas. The herd size was categorized into small (>=15 animals), medium (between 16 and 30 animals), and large (> 30 animals) [25]. Based on parity, cattle were grouped into: no parity (male animals and heifers), 1-3 parity (animals which gave birth up to 3 times) and > 3 (animals which gave birth greater than three). The individual animal was classified as young if it was under 24 months old and as adults if it was equal to or greater than 24 months old.

### Sample size determination

The sample size of the cattle was calculated by the formula described by Thrusfield [26] using an acceptable error of 5% and at a 95% confidence interval. As there is no reported seroprevalence of brucellosis in the study areas, a 50% predicted prevalence and a 95% degree of confidence was employed. Accordingly, the calculated sample size was 384.

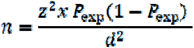

Where n=required sample size

Z=reliability coefficient (1.96 at d=0.05 or 95% CI)

Pexp=expected prevalence (50%)

d= desired absolute precision (95% CI)

For a questionnaire survey, the sample size was calculated using the formula given by Arsham [27], which is as follows:

N = 0.25/SE2,

Where N = sample size and SE (standard error) = 5%.

Thus, the calculated sample size was 100, but 10% of the calculated sample size was added to compensate for non-response rates, which makes the total sample 110.

### Sampling Technique

A multi-stage sampling technique was used to collect representative samples. The Gambella region is divided into three zones, and each zone is divided into districts. Each district is grouped into kebeles, and each kebele is also categorized into different peasant associations or also calledvillage. Accordingly, two districts, Jikawo district and Lare district, were selected purposively based on accessibility and a high cattle population.the two kebeles, eight Villages and households/herds from each district were selected using simple random sampling. Study animals were also selected using simple random sampling. The study animals were stratified according to their age and sex. From each stratum, animals were selected proportionally. Animals below six months of age were excluded from sampling. The total number of samples required was distributed according to the animal population proportionality for each administrative category. The total number of cattle in Jikawo district and Lare district is 38,834 and 45, 540m respectively. A total of 229 and 155 cattle from the Lare district and the Jikawo district, respectively, were considered as study animals. A total of 5,864 households in Jikawo and 5,432 households in Lare are found. For the questionnaire survey, 40 households from the Lare district and 30 from the Jikawo district were randomly selected and considered.

### Sample collection and interview data

Age, sex, herd size, parity, presence of other species, history of abortion and retained fetal membrane were recorded by interviewing the animal attendants or owners while collecting samples. From each study animal, about ten milliliters of blood was aseptically collected from the jugular vein using plain vacutainer tubes and sterile needles. After collection, each vacutainer tube that had a blood sample was placed in an upright position at room temperature for 10 hours to obtain a serum sample. Then sera were decanted into cryovials and labeled. The serum samples were placed in an icebox and transported to the National Animal Health Diagnostic and Investigation Center (NAHDIC), Sebeta, Ethiopia, and kept in a refrigerator at -20 °C until laboratory examination conducted.

### Questionnaire Survey

A pretested KAP questionnaire survey consisting of 30 questions was prepared as the data collection tool. It was divided into four sections: (1) socio-economic characteristic of respondents (2) knowledge of brucellosis (3) attitudes toward brucellosis and (4) practices relating to cattle husbandry, disposal of aborted material and dairy product consumption. The questionnaire survey was closed ended and contained binary and multiple choices. Cattle owners aged at least 15 years, residents in selected kebeles and able to communicate verbally in the local Nyuer language were interviewed face to-face. Cattle owners were randomly selected for a questionnaire survey.

### Serological Tests

#### Rose Bengal Plate Test

All serum samples collected were screened using RBPT according to the procedures described by the World Organization for Animal Health (OIE, 2004) and manufacturers’ instructions. The serum samples were screened using the RBPT antigen (VLA Weybridge, UK). The test serum and antigen were kept at room temperature for half an hour before the test. Then equal volumes (30 μl each) of RBPT antigen and test serum were placed alongside the plate and mixed thoroughly on the clean plate. Both certified reference positive and negative sera were used in each plate for the quality assurance of the result The plate was manually rocked and rotated for 4 minutes, and the degrees of agglutination reactions were recorded. The result was interpreted as Negative if no agglutination and rimming were observed. If barely perceptible agglutination and/or some rimming was considered as 1+ a positive sample, fine agglutination, definite rimming was considered as 2+ positive and clear clumping with definite clearing was considered as 3+ positive

#### Complement Fixation Test (CFT)

A serum sample tested positive by the RBPT was further tested using CFT for confirmation using the standard *B. abortus* antigen (Cenogenics Corporation, USA). The standard *B. abortus* antigen was used to detect the presence of anti-Brucella antibodies in a serum sample. Preparation of the reagent was evaluated by titration and performed according to protocols recommended by World Organization for Animal Health [28]. A certified positive and negative control sera were run together with the samples on each plate as a quality control of the test. A serum sample with a strong reaction, more than 75% fixation of complement (3+) at a dilution of 1:5 or at least with 50% fixation of complement (2+) at a dilution of 1:10, was classified as positive. If there was a lack of fixation or complete hemolysis, it was considered a negative.

### Data management and analysis

The data from the laboratory investigation and the questionnaire survey were entered into a Microsoft Excel spreadsheet, coded, and analyzed with STATA version 14.0 software (Stata Corp, College Station, USA). For the questionnaire survey, descriptive statistics were used to describe the study variables. The overall score was obtained by summing responses from each question and categorizing them into groups, i.e., correct responses 50% to indicate low level, 50-75% correct responses to indicate medium level, and > 75% correct responses to indicate high level for knowledge, practice, and attitude. The seroprevalence of brucellosis was calculated as the number of seropositive samples divided by the total number of samples tested. Similarly, the herd level prevalence was calculated by dividing the number of herds with at least one animal positive for brucellosis by the total number of herds tested [29]. Descriptive statistics were used to summarize seroprevalence, whereas logistic regression was used to assess the association of risk factors with seroprevalence of Brucella antibodies. Potential risk factors considered for statistical analysis include age, sex, parity, herd size, abortion history, presence of other species, and district. For all risk factors, the level with the lowest prevalence was used as a reference category. All variables having a p-value of <0.25 in the univariable logistic regression analysis were further analyzed by multivariable logistic regression after checking for co-founders. In all the tested variables, p<0.05 was set for significance, and the variables with p<0.05 in the multivariable model were concluded as predicting factors for seropositivity of brucellosis.

## Results

### Serological Analysis

A total of 384 sera samples were collected from 70 herds of cattle and screened with RBPT and confirmed with CFT. Out of 384 serum samples, 6.77% (26/384) and 3.13% (12/384) were found to be RBPT positive and CFT positive, respectively, at the animal level. The CFT result showed that Jikawo district had a higher seroprevalence of bovine brucellosis at both the individual animal (5.16%) and herd level (16.67%) than Lare district (Table 1).

**Table 1.** Seroprevalence of bovine brucellosis in Lare and Jikawo districts, Gambella, Ethiopia

| Variable | Animal level |  |  | Herd level |  |  |
| --- | --- | --- | --- | --- | --- | --- |
| Study area | No of animal examined | RBPT | CFT | No of herd examined | RBPT | CFT |
| Lare | 229 | 12(5.24) | 4(1.75) | 40 | 9(22.50) | 4(10) |
| Jikawo | 155 | 14(9.03) | 8(5.16) | 30 | 8(26.67) | 5(16.67) |
| Total | 384 | 26(6.77) | 12(3.13) | 70 | 17(24.29) | 9(12.85) |

The univariable logistic regression analysis showed that the risk of bovine brucellosis in the Jikawo district is 3.06 times higher than in the Lare district. Adult cattle are more likely to be affected by brucellosis (OR = 4.39) than young cattle. Similarly, cattle kept mixed with small ruminants had a higher probability of being infected by brucellosis than cattle kept alone (OR = 4.35). The probability of being affected by brucellosis was also significantly higher in cattle with a parity of greater than three as compared to cattle with no parity. The multicollinearity matrix result revealed that all independent variables were not collinear with each other (r<0.5) except parity versus age (r=0.66), abortion history versus age (r=.71) and abortion history versus parity (r=0.91). Thus, considering univariable p-value < 0.25, non-collinearity, and frequency of variable categories, the following variables were selected for entry into the multivariable model: district, herd size, and presence of other species. The multivariable logistic regression model revealed that herd size (OR= 4.65; 95% CI: 1.63-13.28, p<0.05) and presence of other species (OR= 4.91; 95% CI: 1.02-23.76, p<0.05) were potential risk factors for cattle seropositivity to circulating Brucella antibodies and independent predictors of bovine brucellosis in the study area (Table 2).

**Table 2.** Univariable and multivariable logistic regression analysis of risk factors for Brucella seropositivity

| Risk factors | Category | No. Exam. | No. Positive (%) | Univariable |  |  | Multivariable |  |  |
| --- | --- | --- | --- | --- | --- | --- | --- | --- | --- |
|  |  |  |  | OR | P-value | 95% CI | OR | P-value | 95% CI |
| Districts | Lare | 229 | 4(1.75%) | 1.0 | - | - | 1.0 | - | - |
|  | Jikawo | 155 | 8(5.16%) | 3.06 | 0.072 | 0.91-10.35 | 2.84 | 0.101 | 0.81-9.91 |
| Herd size | Medium | 111 | 1(0.90%) | 1.0 | - | - | 1.0 | - | - |
|  | Large | 216 | 5(2.31%) | 2.61 | 0.385 | 0.30-22.59 |  |  |  |
|  | Small | 57 | 6(10.53%) | 12.94 | 0.019 | 1.52-110.31 | 4.65 | 0.004 | 1.63-13.28 |
| Sex | Male | 130 | 3(2.31%) | 1.0 | - | - |  |  |  |
|  | Female | 254 | 9(3.59%) | 1.56 | 0.513 | 4.14-5.85 |  |  |  |
| Age | Young | 176 | 2(1.14%) | 1.0 | - | - |  |  |  |
|  | Adult | 208 | 10(4.81%) | 4.39 | 0.058 | 0.95-20.33 |  |  |  |
| Parity | No parity | 238 | 3(1.26%) | 1.0 | - | - |  |  |  |
|  | 1-3 | 71 | 4(5.63%) | 4.68 | 0.047 | 1.02-21.41 |  |  |  |
|  | >3 | 75 | 5(6.67%) | 5.60 | 0.020 | 1.31-23.10 |  |  |  |
| Abortion history | Not concerned | 237 | 3(1.28%) | 1.0 | - | - |  |  |  |
|  | No | 144 | 7(4.79%) | 4.03 | 0.046 | 1.03-15.85 |  |  |  |
|  | Yes | 3 | 2(66.67) | 156.67 | 0.000 | 10.10-2232.78 |  |  |  |
| Presence of other species | No | 175 | 2(1.14%) | 1.0 | - | - | 1.0 | - | - |
|  | Yes | 209 | 10(4.7%) | 4.35 | 0.060 | 0.94-20.11 | 4.91 | 0.048 | 1.02-23.76 |

### Questionnaire Survey Analysis

#### Socio-economic characteristics of respondents

A total of 110 cattle owners were interviewed during the study period, of which 92 (83.6%) were male. The respondents’ educational level showed that most of them (87.3% [96/110]) are illiterate. The income source of most respondents (52.7%) was based on animal sales, followed by animal and dairy product sales (32.7%) as shown in Table 3.

**Table 3.** Socio-economic characteristics of respondents in Lare and Jikawo districts

| Variables | Categories | Frequency | Percent |
| --- | --- | --- | --- |
| Educational level | Illiterate | 96 | 87.3 |
|  | Primary | 13 | 11.8 |
|  | Secondary and above | 1 | 0.9 |
| Age | 21-35 | 35 | 31.8 |
|  | 36-49 | 41 | 37.3 |
|  | >49 | 34 | 30.9 |
| Sex | Female | 18 | 16.4 |
|  | Male | 92 | 83.6 |
| Family size | 3-6 | 44 | 40 |
|  | 7-10 | 42 | 38.2 |
|  | >10 | 24 | 21.8 |
| Source income | Crop sale | 1 | 0.9 |
|  | Animal sale | 58 | 52.7 |
|  | Dairy product sale | 15 | 13.6 |
|  | Animal and dairy product sale | 36 | 32.7 |

### Analysis of knowledge, attitude, and practice of respondents

Mostrespondents (66.4% [73/110]) had heard about brucellosis. However, 92.7% of the respondents (102/110) did not know that brucellosis is a zoonotic disease, 77.2% (86/100) did not know that brucellosis causes abortion, and 89.1% (98/100) did not know that brucellosis can be transmitted to humans by handling aborted fetus and consumption of raw milk from infected cows. As part of the preventive measures of brucellosis adopted by cattle owners, most suggested using boiled milk, while others suggested testing and culling and improved sanitation. A few of them, 11.8% (13/110), never knew any control and preventive measure (Table 4).

**Table 4.**
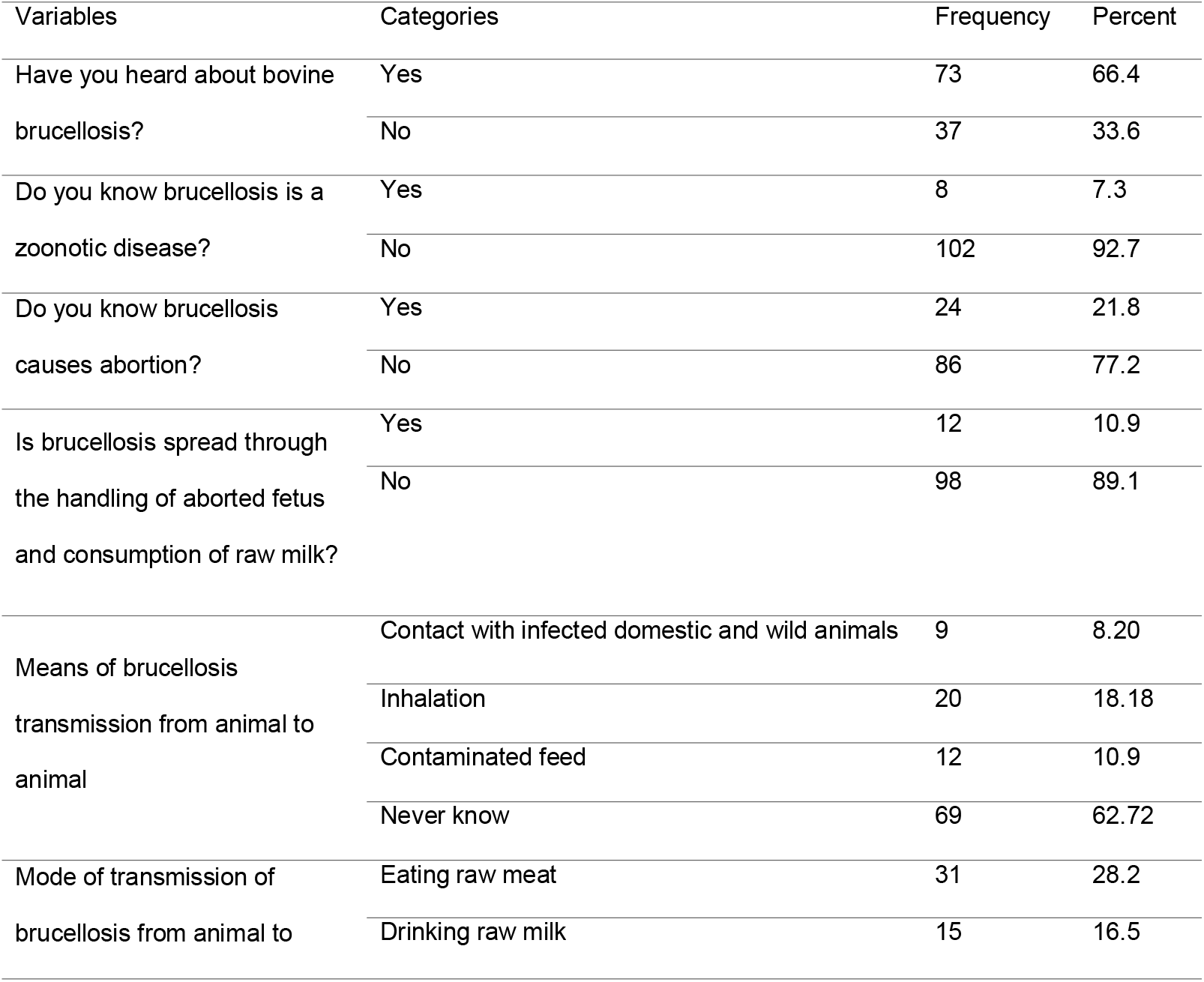
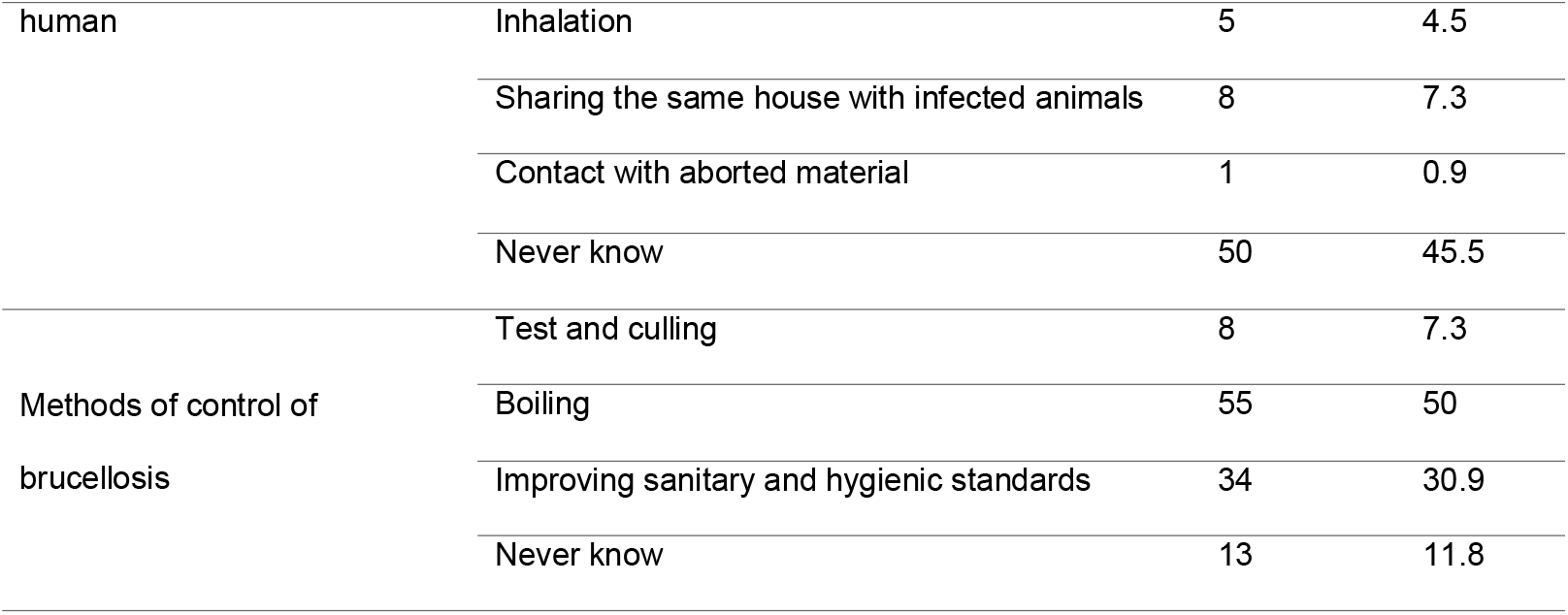
Respondents’ knowledge of brucellosis in the study area

Analysis of the attitude of respondents showed that only 17.3% (19/110) believed that some of their family members were at risk of contracting brucellosis if exposed to infected cattle.

Moreover, most respondents do not think boiling milk before consumption, useinggloves when handling infected cattle or aborted material and washing hands after close contact with infected or aborted material are necessary to prevent transmission of bovine brucellosis to humans (Table 5). Table 5. Attitude of respondents toward brucellosis in study areas

**Table 5.** Attitude of respondents toward brucellosis in study areas

| Variable | Category | Frequency | Percent |
| --- | --- | --- | --- |
| Do you believe infected cattle can expose family<br>members to *Brucella* infection? | Yes | 19 | 17.3 |
|  | No | 91 | 82.7 |
| Do you think boiling milk is necessary before<br>consumption to prevent brucellosis? | Yes | 39 | 35.5 |
|  | No | 71 | 64.5 |
| Do you think it is necessary to use gloves when<br>handling infected cattle or aborted material? | Yes | 14 | 12.7 |
|  | No | 96 | 87.3 |
| Do you think washing your hands is necessary after<br>close contact with animals or their abortus? | Yes | 25 | 22.7 |
|  | No | 85 | 77.3 |
| Do you think the use of vaccination is necessary to<br>prevent brucellosis? | Yes | 109 | 99.1 |
|  | No | 1 | 0.9 |

Most respondents used to practice risky activities such as not washing of hands before and after milking (80%), disposing of an aborted fetus with bare hands (90%), disposing of an aborted fetus in open fields (82.2%), handling animals with uncovered wounds (100%), and consumption of raw milk (85.5%) (Table 6).

**Table 6.** Practices of respondents regarding bovine brucellosis in the study areas

| Variable | Category | Frequency | Percent |
| --- | --- | --- | --- |
| What type of Housing system do you use? | Open field | 110 | 100 |
| Do you practice hand washing before and after milking | Yes | 22 | 20 |
|  | No | 88 | 80 |
| How do you dispose aborted fetus? | By protective materials | 11 | 10 |
|  | By uncovered hand | 99 | 90 |
| Where do you dispose aborted fetus? | Incineration | 4 | 3.6 |
|  | Deep burial | 6 | 5.5 |
|  | Disposing to open field | 91 | 82.2 |
|  | Throw it away for carnivores | 9 | 8.2 |
| Do you keep other animals in the herd? | Yes | 56 | 49.1 |
|  | No | 54 | 50.9 |
| Do you use of protective materials during assisting parturition? | Yes | 17 | 15.5 |
|  | No | 93 | 84.5 |
| Do you covering wound while handling animals? | Yes | 10 | 9.1 |
|  | No | 100 | 90.9 |
| Form of milk Consumed? | Raw | 94 | 85.5 |
|  | Boiled | 9 | 8.9 |
|  | Processed | 7 | 6.4 |
| Do you assist dairy cows during parturition? | Yes | 89 | 80.9 |
|  | No | 21 | 19.1 |
| Do you consumption of raw milk? | Yes | 104 | 94.54 |
|  | No | 6 | 5.46 |
| How do you dispose animals died of suspected of brucellosis? | Burn carcass | 9 | 8.18 |
|  | Burring all carcass | 7 | 6.36 |
|  | Cook and eat the meat | 2 | 1.82 |
|  | Disposing to open field | 92 | 83.64 |

## Discussion

The present study revealed that the overall seroprevalence of bovine brucellosis was 3.13% in the Lare and Jikawo districts of the Gambella region at the individual animal level. This value was consistent with the 2.9% prevalence in Asella by Tsegaye et al. [30] and 3.19% in the Tigray region by Berhe et al. [31]. However, the current prevalence was higher than the previous reports of Degefu et al. [32] 1.38% in Jijjiga Zone, Somalia, Kassahun et al. [33] 1.92% in Sidama Zone, Yohannes et al. [34] 1.97% in Guto-Gida district of East Wollega Zone, Bashitu et al. [35] 0.2% in Ambo and 0% in Debrebirhan town. In contrast to the current finding, higher seroprevalence was reported by Haileselassie et al. [36] at 7.7% in the Tigray region, Ibrahim et al. [15] at 15.0% in the Jimma zone of the Oromia region, Dinka and Chala [37] at 11.2% in the East Shewa Zone of the Oromia region, and Berhe et al. [31] at 42.3% in the extensive cattle production system of the Tigray Region of Ethiopia. Also, a higher prevalence than in the current study has been reported in different African countries, such as Ivory Coast (8.8–10.3%) by Sanogo et al. [38], Zambia (18.7%) by Chimana et al. [39], and Algeria (9.7%) by Aggad and Boukraa [40]. The variation in prevalence reported from different regions of Ethiopia and other parts of Africa could be associated with the evolution of the disease, geographical origin, breeds, sample size, cattle rearing system, study frame, as well as the protocol adopted, such as the type and number of diagnostic tests used. The series of serology protocols used by researchers to screen and to confirm the disease might be one test or more than one test, i.e., a screening test followed by confirmation of positive reactors by another test or in parallel interpretation [41].

The present study showed no statistically significant difference in the seroprevalence of brucellosis between the two districts (Lare and Jikawo). This could be due to the similarity of traditional cattle management systems in both districts where pastoral livestock raising is predominant [15, [31, [42]. In the current study, there was a higher seroprevalence of brucellosis in adult cattle than in young cattle. This finding agrees with the reports of Kassahun et al. [33] and Adugna et al. [42]. It has also been well documented that brucellosis is more associated with sexual maturity [43], and a higher seroprevalence has been repeatedly reported in sexually matured animals.

The present study revealed that the presence of other livestock (sheep or goats) was the risk factor associated with the presence of seroreactor cattle. Although sheep and goats were not tested for brucellosis in this study, the finding corroborates reports of mixed farming importance in Brucella transmission dynamics in Egypt [44]. On the other hand, *B. abortus* infection was isolated and reported from sheep and goats in Nigeria by Ocholi et al. [45], and *B. melitensis* was isolated from cattle in Egypt by Samaha et al. [44]. Accordingly, contact between cattle and sheep and goats was the most important risk factor identified in these studies. Thus, segregating sheep and goats from cattle may reduce the seroprevalence and spread of B. melitensis among cattle in mixed herds.

The prevalence of brucellosis was significant in cows with a history of abortion in the current study. Different authors also reported a different prevalence of brucellosis in cattle with history of abortion [15, [30, [31, [42, [46]. The female animals were more positive reactors than male animals in this study. It has been reported that males are usually more resistant than female cattle [31, [46, [47, [48]. Different factors are probably involved in the variation in sex susceptibility, including physiological and behavioral differences between males and females. Because of the preferential growth of *B. abortus* in the gravid uterus, it can enter the uterus as it disseminates from the main sites of carrier states (udder, supramammary lymph node) [43].

The existence of a previous history of abortion was significantly associated with the prevalence of brucellosis (p<0.05) in the present study. This finding is in agreement with some of studies, where significant associations between seropositivity and history of abortion have been reported [15, [42, [46, [49]. Similarly, studies in different African countries also show that individual animal brucellosis seroprevalence correlates with the presence of abortions [27, [48, [50]. This could be explained by the fact that abortion is a typical outcome of brucellosis [49, [51, [52].

Based on parity, the difference observed in seroprevalence was statistically insignificant. Similar observations were recorded by Minda et al. [51] and Berhe et al. [31]. Although there is an insignificant association between parity and brucellosis seropositivity, a higher seroprevalence was observed in cattle with greater than three parturition (6.67%) than in cattle with one up to three parturition (5.63%) in the study area. The higher seroprevalence of brucellosis in the multi-parturition cattle of this study was in line with the findings of Minda et al. [51] and Asmare et al. [53].

Improvement of knowledge, attitudes, and practices among cattle owners could have a significantly impact the reduction of many zoonotic infections, including brucellosis. The analysis of the KAP in the current study showed that most cattle owners in the studied area had heard about bovine brucellosis (66.4%), but most respondents did not know it was a zoonotic disease (92.7%). Similar results were reported in brucellosis KAP studies conducted in northern Uganda [54], and Kenya [55] where 63% and 79% of community participants had heard of brucellosis, respectively. Studies conducted in Egypt, [56] Nigeria, [57] Uganda, [58] and Jordan [59] showed that 83%, 93%, 99.3%, and 100% had heard of brucellosis, respectively. Contrasting results were found in a brucellosis KAP study in Tajikistan, where only 15% had heard of brucellosis [60]. The primary sources of brucellosis information were stated as unspecified media in the Jordan study, [59] community health workers in the Kenya study, [55] parents in the Nigeria study, [57] and friends or family members in the Tajikistan study [60]. Most of the respondents had heard about brucellosis from veterinarians working in veterinary clinics, indicating the importance of the role of government veterinary services in the current study. Poor hygienic practices and uncontrolled animal movements were practiced in extensive husbandry systems. This could pose a substantial risk of transmitting the disease within and in between the herds. The present study findings also agree with previous studies on the intensive farming system in Ethiopia [51, [61, [62, [63, [64].

Cattle owners’ knowledge, attitude, and practice regarding the disease are crucial steps in developing prevention and control measures [65]. In the current study, most respondents have limited knowledge and attitudes about disease transmission and control. Moreover, they have been practicing risky activities such as assisting their animals during parturition, disposing of aborted fetuses and afterbirth in an open environment without protective gloves or masks, and consuming raw milk. These might have resulted in high risks of disease transmission within and between the herds and humans. The current findings agrees with previous studies on extensive livestock production system [25, [42, [66]. The occurrence of brucellosis in humans is associated with contacting aborted animals with bare hands and assisting animals during parturition [67].

## Conclusion

The present study revealed a 3.13% and 12.5% overall seroprevalence of bovine brucellosis at individual animal and herd levels, respectively, in the Gambella region, Ethiopia. The seroprevalence of the disease was associated with the presence of small ruminants and size of the cattle herd. The present study also found that cattle owners’ knowledge, attitude, and practice toward brucellosis in the study were low. This might contribute to the wide spread of bovine brucellosis both in animals and humans. Therefore, creating awareness for the community on the mechanisms of transmission, zoonotic importance, prevention, control, and economic importance of the disease is recommended. Moreover, communication and cooperation between animal and human health professionals, the agricultural and education sectors, cattle owners, and other relevant stakeholders need to be strengthened to reduce disease transmission between animals and human and improve control of the brucellosis.

## Ethics approval and consent to participate

This study was conducted following the Declaration of Helsinki. All study animal owners were informed about the study and informed consents were obtained from all cow owners and individuals participated in this study. Participation in the study was on voluntary bases. Confidentiality was assured by using codes. We confirm that the animals were handled with the best practices of veterinary care. Ethical clearance was obtained from the Ambo University research and ethical review committee (Ref.No. □ □/42/31/40/12).

## Consent for publication

Not applicable.

## Data Availability Statement

The data generated and analyzed during the current study are available in the (raw data compiled.xls) deposited in Open Science Framework (OSF) repository as (https://osf.io/g826x/files/osfstorage/63d31c938a2ec2010e635187).

## Acknowledgments

The authors thank the NAHDIC for their unreserved laboratory facilities and reagent support and the staff members for their technical support in the laboratory analysis. We acknowledge the animal owners and animal health assistants of the different districts for their cooperation during sample collection.

## References

1. Akinseye VO, Adesokan HK, Ogugua AJ, Adedoyin FJ, Otu PI, Kwaghe AV, et al. Sero-epidemiological survey and risk factors associated with bovine brucellosis among slaughtered cattle in Nigeria. Onderstepoort J Vet Res. 2016;83(1):a1002.

2. McDermott J, Grace D, Zinsstag J. Economics of brucellosis impact and control in low-income countries. Rev Sci Tech. 2013;32(1):249–61.

3. Angesom H, Mahendra P, Tesfu K, Fikre Z. Sero-epidemiology of camel brucellosis in the Afar region of Northeast Ethiopia. Journal of Veterinary Medicine and Animal Health. 2013;5(9):269–75.

4. OIE. Brucellosis: B. abortus, B. melitensis and B. suis. Manual of diagnostic tests and vaccines for terrestrial animals, Paris. 2016. p. 1–44.

5. Onunkwo J, Njoga E, Nwanta J, Shoyinka S, Onyenwe I, Eze J. Serological survey of porcine Brucella infection in Southeast, Nigeria. Nigerian Veterinary Journal. 2011;32(1).

6. Kebede T, Ejeta G, Ameni G. Seroprevalence of bovine brucellosis in smallholder farms in central Ethiopia (Wuchale-Jida district). Revue de Médecine Vétérinaire. 2008;159(1):3.

7. Al-Majali AM, Talafha AQ, Ababneh MM, Ababneh MM. Seroprevalence and risk factors for bovine brucellosis in Jordan. J Vet Sci. 2009;10(1):61–5.

8. Mugizi DR, Boqvist S, Nasinyama GW, Waiswa C, Ikwap K, Rock K, et al. Prevalence of and factors associated with Brucella sero-positivity in cattle in urban and peri-urban Gulu and Soroti towns of Uganda. J Vet Med Sci. 2015;77(5):557–64.

9. Yohannes Gemechu M, Paul Singh Gill J. Seroepidemiological survey of human brucellosis in and around Ludhiana, India. Emerging Health Threats Journal. 2011;4(1):7361.

10. Mukhtar F, Kokab F. Brucella serology in abattoir workers. J Ayub Med Coll Abbottabad. 2008;20(3):57–61.

11. Bechtol D, Carpenter LR, Mosites E, Smalley D, Dunn JR. Brucella melitensis infection following military duty in Iraq. Zoonoses Public Health. 2011;58(7):489–92.

12. Boukary AR, Saegerman C, Abatih E, Fretin D, Alambédji Bada R, De Deken R, et al. Seroprevalence and potential risk factors for Brucella spp. infection in traditional cattle, sheep and goats reared in urban, periurban and rural areas of Niger. PLoS One. 2013;8(12):e83175.

13. Zeng J, Duoji C, Yuan Z, Yuzhen S, Fan W, Tian L, et al. Seroprevalence and risk factors for bovine brucellosis in domestic yaks (Bos grunniens) in Tibet, China. Trop Anim Health Prod. 2017;49(7):1339–44.

14. Al Dahouk S, Neubauer H, Hensel A, Schöneberg I, Nöckler K, Alpers K, et al. Changing epidemiology of human brucellosis, Germany, 1962-2005. Emerg Infect Dis. 2007;13(12):1895–900.

15. Ibrahim N, Belihu K, Lobago F, Bekana M. Sero-prevalence of bovine brucellosis and its risk factors in Jimma zone of Oromia Region, South-western Ethiopia. Trop Anim Health Prod. 2010;42(1):35–40.

16. Hailemelekot M, Kassa T, Tefera M, Belihu K, Asfaw Y, Ali A. Seroprevalence of brucellosis in cattle and occupationally related humans in selected sites of Ethiopia. Ethiopian Veterinary Journal. 2007;11:85–100.

17. Asmare K, Krontveit RI, Ayelet G, Sibhat B, Godfroid J, Skjerve E. Meta-analysis of Brucella seroprevalence in dairy cattle of Ethiopia. Trop Anim Health Prod. 2014;46(8):1341–50.

18. Tadesse G. Brucellosis Seropositivity in Animals and Humans in Ethiopia: A Meta-analysis. PLoS Negl Trop Dis. 2016;10(10):e0005006.

19. Smits HL, Kadri SM. Brucellosis in India: a deceptive infectious disease. Indian J Med Res. 2005;122(5):375–84.

20. Roth F, Zinsstag J, Orkhon D, Chimed-Ochir G, Hutton G, Cosivi O, et al. Human health benefits from livestock vaccination for brucellosis: case study. Bulletin of the World health Organization. 2003;81:867–76.

21. Pieracci EG, Hall AJ, Gharpure R, Haile A, Walelign E, Deressa A, et al. Prioritizing zoonotic diseases in Ethiopia using a one health approach. One Health. 2016;2:131–5.

22. CSA. Central Statistics Authority of Ethiopia Agricultural Sample Survey. 2011.

23. CSA. Statistical Report in characterization of Agricultural household and land use, Part I Addis Ababa, Ethiopia. 2008.

24. Dika G. Impacts of climate variability and households adaptation strategies in lare district of Gambella region, South Western Ethiopia. Journal of Earth Science & Climatic Change. 2018;9(2):7.

25. Megersa B, Biffa D, Niguse F, Rufael T, Asmare K, Skjerve E. Cattle brucellosis in traditional livestock husbandry practice in Southern and Eastern Ethiopia, and its zoonotic implication. Acta Vet Scand. 2011;53(1):24.

26. Thrusfield M. Veterinary Epidemiology. 3rd Edition ed: Wiley; 2007.

27. Arsham H. Questionnaire Design and Surveys Sampling Error. Hyperlink Reference not Valid. 2002.

28. OIE. Terrestrial Animal Health Code Brucellosis, science and Comparative Medicine. 2009. p. 69–98.

29. Alehegn E, Tesfaye S, Chane M. Seroprevalence of Bovine Brucellosis and its risk factors in cattle in and around Gondar Town, North West Gondar, Ethiopia. Journal of Advanced Dairy Research. 2017;4:166.

30. Tsegaye Y, Kyule M, Lobago F. Seroprevalence and risk factors of bovine brucellosis in Arsi Zone, Oromia Regional State, Ethiopia. American Academic Scientific Research Journal for Engineering, Technology, and Sciences. 2016;24(1):16–25.

31. Berhe G, Belihu K, Asfaw Y. Seroepidemiological investigation of bovine brucellosis in the extensive cattle production system of Tigray region of Ethiopia. International Journal of Applied Research in Veterinary Medicine. 2007;5(2):65.

32. Degefu H, Mohamud M, Hailemelekot M, Yohannes M. Seroprevalence of bovine brucellosis in agro pastoral areas of Jijjiga zone of Somali National Regional State, Eastern Ethiopia. Ethiopian Veterinary Journal. 2011;15(1).

33. Kassahun A, Yilkal A, Esayas G, Gelagay A. Brucellosis in extensive management system of Zebu cattle in Sidama Zone, Southern Ethiopia. African Journal of Agricultural Research. 2010;5(3):257–63.

34. Yohannes M, Degefu H, Tolosa T, Belihu K, Cutler RR, Cutler S. Brucellosis in Ethiopia. African Journal of Microbiology Research. 2013;7(14):1150–7.

35. Bashitu L, Afera B, Tuli G, Aklilu F. Sero-prevalence study of bovine brucellosis and its associated risk factors in Debrebirhan and Ambo towns. J Adv Dairy Res. 2015;3(131):2.

36. Haileselassie M, Shewit K, Moses K. Serological survey of bovine brucellosis in barka and arado breeds (Bos indicus) of western Tigray, Ethiopia. Prev Vet Med. 2010;94(1-2):28–35.

37. Dinka H, Chala R. Seroprevalence study of bovine brucellosis in pastoral and agro-pastoral areas of East Showa Zone, Oromia Regional State, Ethiopia. American-Eurasian Journal of Agricultural and Environmental Science. 2009;6(5):508–12.

38. Sanogo M, Abatih E, Thys E, Fretin D, Berkvens D, Saegerman C. Risk factors associated with brucellosis seropositivity among cattle in the central savannah-forest area of Ivory Coast. Prev Vet Med. 2012;107(1-2):51–6.

39. Chimana HM, Muma JB, Samui KL, Hangombe BM, Munyeme M, Matope G, et al. A comparative study of the seroprevalence of brucellosis in commercial and small-scale mixed dairy-beef cattle enterprises of Lusaka province and Chibombo district, Zambia. Trop Anim Health Prod. 2010;42(7):1541–5.

40. Aggad H, Boukraa L. Prevalence of bovine and human brucellosis in western Algeria: comparison of screening tests. East Mediterr Health J. 2006;12(1-2):119–28.

41. Chisi SL, Marageni Y, Naidoo P, Zulu G, Akol GW, Van Heerden H. An evaluation of serological tests in the diagnosis of bovine brucellosis in naturally infected cattle in KwaZulu-Natal province in South Africa. J S Afr Vet Assoc. 2017;88(0):e1–e7.

42. Adugna KE, Agga GE, Zewde G. Seroepidemiological survey of bovine brucellosis in cattle under a traditional production system in western Ethiopia. Rev Sci Tech. 2013;32(3):765–73.

43. Radostits OM, Done SH. Veterinary Medicine: A Textbook of the Diseases of Cattle, Sheep, Pigs, Goats, and Horses: Elsevier Saunders; 2007.

44. Samaha H, Al-Rowaily M, Khoudair RM, Ashour HM. Multicenter study of brucellosis in Egypt. Emerging infectious diseases. 2008;14(12):1916.

45. Ocholi RA, Kwaga JK, Ajogi I, Bale JO. Phenotypic characterization of Brucella strains isolated from livestock in Nigeria. Vet Microbiol. 2004;103(1-2):47–53.

46. Tolosa T, Regassa F, Belihu K. Seroprevalence study of bovine brucellosis in extensive management system in selected sites of Jimma Zone, Western Ethiopia. Bulletin of Animal health and Production in Africa. 2008;56(1).

47. Bayemi PH, Webb EC, Nsongka MV, Unger H, Njakoi H. Prevalence of Brucella abortus antibodies in serum of Holstein cattle in Cameroon. Trop Anim Health Prod. 2009;41(2):141–4.

48. Muma JB, Pandey GS, Munyeme M, Mumba C, Mkandawire E, Chimana HM. Brucellosis among smallholder cattle farmers in Zambia: public health significance. Trop Anim Health Prod. 2012;44(4):915–20.

49. Alemu F, Admasu P, Feyera T, Niguse A. Seroprevalence of bovine brucellosis in eastern Showa, Ethiopia. Academic Journal of Animal Diseases. 2014;3(3):27–32.

50. Schelling E, Diguimbaye C, Daoud S, Nicolet J, Boerlin P, Tanner M, et al. Brucellosis and Q-fever seroprevalences of nomadic pastoralists and their livestock in Chad. Prev Vet Med. 2003;61(4):279–93.

51. Minda AG, Gobena A, Tesfu k, Getachew T, Angella A, Gezahegne MK. Seropositivity and risk factors for Brucella in dairy cows in Asella and Bishoftu towns, Oromia Regional State, Ethiopia. African Journal of Microbiology Research. 2016;10(7):203–13.

52. Eyob E, Hani S, Diriba L, Birhanu A. Prevalence and risk analysis of bovine brucellosis in Asella organized dairy farm, Oromia Regional State, South East Ethiopia. Journal of Veterinary Medicine and Animal Health. 2018;10(10):245–9.

53. Asmare K, Sibhat B, Molla W, Ayelet G, Shiferaw J, Martin AD, et al. The status of bovine brucellosis in Ethiopia with special emphasis on exotic and cross bred cattle in dairy and breeding farms. Acta Trop. 2013;126(3):186–92.

54. Nabirye HM, Erume J, Nasinyama GW, Kungu JM, Nakavuma J, Ongeng D, et al. Brucellosis: Community, medical and veterinary workers’ knowledge, attitudes, and practices in Northern Uganda. 2017.

55. Obonyo M, Gufu WB. Knowledge, attitude and practices towards brucellosis among pastoral community in Kenya, 2013. International Journal of Innovative Research and Development. 2015;4(10):375–84.

56. Holt HR, Eltholth MM, Hegazy YM, El-Tras WF, Tayel AA, Guitian J. Brucella spp. infection in large ruminants in an endemic area of Egypt: cross-sectional study investigating seroprevalence, risk factors and livestock owner’s knowledge, attitudes and practices (KAPs). BMC Public Health. 2011;11:341.

57. Buhari H, Saidu S, Mohammed G, Raji M. Knowledge, attitude and practices of pastoralists on bovine brucellosis in the north senatorial district of Kaduna state, Nigeria. J Anim Health Prod. 2015;3(2):28–34.

58. Kansiime C, Mugisha A, Makumbi F, Mugisha S, Rwego IB, Sempa J, et al. Knowledge and perceptions of brucellosis in the pastoral communities adjacent to Lake Mburo National Park, Uganda. BMC Public Health. 2014;14:242.

59. Musallam, II, Abo-Shehada MN, Guitian J. Knowledge, Attitudes, and Practices Associated with Brucellosis in Livestock Owners in Jordan. Am J Trop Med Hyg. 2015;93(6):1148–55.

60. Lindahl E, Sattorov N, Boqvist S, Magnusson U. A study of knowledge, attitudes and practices relating to brucellosis among small-scale dairy farmers in an urban and peri-urban area of Tajikistan. PLoS One. 2015;10(2):e0117318.

61. Tesfaye G, Tsegaye W, Chanie M, Abinet F. Seroprevalence and associated risk factors of bovine brucellosis in Addis Ababa dairy farms. Trop Anim Health Prod. 2011;43(5):1001–5.

62. Asgedom H, Damena D, Duguma R. Seroprevalence of bovine brucellosis and associated risk factors in and around Alage district, Ethiopia. Springerplus. 2016;5(1):851.

63. Elemo KK, Geresu MA. Bovine brucellosis: Seroprevalence and its associated risk factors in cattle from smallholder farms in Agarfa and Berbere districts of Bale Zone, South Eastern Ethiopia. Journal of Animal and Plant Sciences. 2018;28(2):28–2.

64. Waktole H, Geneti E, Ahmed WM, Mammo G, Abunna F. Sero-Prevalence and Associated Risk Factors of Bovine Brucellosis in Selected Dairy Farms in Bishoftu Town, Oromia, Ethiopia. Intl J. 2018;9(2):45–53.

65. Prilutski MA. A brief look at effective health communication strategies in Ghana. Elon J Undergrad Res Commun. 2010;1:51–8.

66. Genene R, Desalew M, Yamuah L, Hiwot T, Teshome G, Asfawesen G, et al. Human brucellosis in traditional pastoral communities in Ethiopia. International Journal of Tropical Medicine. 2009;4(2):59–64.

67. Kozukeev T, Maes E, Favorov M. Centers for Disease Control and Prevention (CDC) Risk factors for brucellosis—Leylek and Kadamjay districts, Batken Oblast, Kyrgyzstan. MMWR Suppl. 2006;1:31–4.

